# Expression of Non-Visual Opsins in the Green Anole Lizard (*Anolis carolinensis*)

**DOI:** 10.1101/2025.03.20.644269

**Authors:** Violeta Trejo-Reveles, Grace E. Anderson, Alex R. Johnston, Simone L. Meddle, Michele A. Johnson

## Abstract

Extra-retinal photoreception is widely observed across vertebrates, facilitating species-specific regulation of physiological and behavioral responses to diverse photic environments. Yet, the roles of non-visual opsins remain poorly understood, as their distribution and function have not yet been systematically investigated. This study provides the first comprehensive evidence of non-visual opsin expression across the reptile body. Using predicted sequences for extra-retinal photoreceptors, non-visual opsins were confirmed in the brain, eye, testes, liver, and skin. Specifically, OPN3 (encephalopsin) exhibited the highest expression across all tissues, particularly in the testes, corroborating findings in mammals. OPN5 (neuropsin) was detected predominantly in the testes, marking the first such report in reptiles. Moderate expression of OPN4 (melanopsin) in the brain and eye supports its roles in both visual and deep-brain photoreception. In the skin, a distinct pattern of higher opsin expression was observed in the dorsal skin compared to ventral skin. OPN3-3 was most abundant and exhibited a unique expression profile, potentially linking opsins to light-mediated processes influencing chromatophore-driven color changes. Findings from this research provide a critical evolutionary context for understanding the role and conservation of non-visual opsins in reptiles and their relevance to vertebrate lineages, including mammals and birds.

## Introduction

Extra-retinal photoreception, mediated by photosensitive molecules known as opsins, is prevalent across vertebrates. Decades of investigation have unveiled distinct families of non-visual opsins. These extra-retinal photoreceptors (ERPs) play a pivotal role in orchestrating a myriad of physiological, metabolic, behavioral, and morphological responses to light stimuli, yet the precise mechanisms governing fluctuations in these light-mediated traits remain largely unknown [35]. ERPs are typically comprised of a photosensitive opsin protein and a chromophore, commonly an 11-cis-retinal, although all-trans-retinal chromophores are also present. These non-visual photoreceptors utilize G protein-coupled transduction pathways for signaling, but unlike visual opsins, most non-visual opsins form bi-stable pigments that transition between light and dark states. This happens solely through light exposure, without external input to return to the dark state [32]. Structural variations in opsins result in a diverse range of absorption spectra, most likely reflecting adaptations to species-specific photic environments. Recent studies have identified opsin localization in the brain of vertebrate species. In the brain, ERPs play an important role in mediating various body functions including migration, circadian rhythms, skin color change, and reproduction [35]. Most opsins have shown high expression in the hypothalamus [15] and the pineal gland [28, 2, 4], but there is also evidence to suggest that ERPs are expressed in the cerebellum and preoptic area regions of the brain [5, 6]. Opsins are also found in other internal organs such as the testes, liver, and kidneys; however, the role they play within these tissues is currently unclear [37, 47].

Studies on non-visual opsins in mammals, birds, fish, and frogs have emerged in the past decade, but remarkably little is known regarding the distribution or function of non-visual opsins in non-avian reptiles (hereafter, reptiles). Yet, reptiles occupy a position of critical importance in the amniote phylogeny, as the reptilian clade (Sauropsida) includes birds, and this group is the closest relative of mammals. Thus, studies of reptile opsins can provide the evolutionary context to understand variation in opsins in the avian and mammalian lineages. In lizards, most opsin studies to date have focused on visual opsin expression [49, 34, 48]. For example, one of the first studies to examine extra-retinal photoreception in lizards was Underwood and Menaker in 1976 [49]. The authors reported greater expression of OPNX in lizards exposed to light, but at the time opsin transcripts in lizards were not completely annotated. It was not until 25 years later that brain opsin cDNA, that appeared to belong to the RH2 class of cone-opsins involved in circadian rhythms, was identified in lizards [34]. Other studies focused on retinal and extra-retinal photoreceptor expression in the reptilian eye, such as pinopsin expression in the gecko retina [44] and different visual opsins and sex-based differences in the tawny dragon (*Ctenophorus decresii*) retina [11]. However, there is little information about the roles non-visual opsins play in lizards, whether their expression is conserved, and if they have similar expression patterns to that of mammals and birds. An early study focused on the expression of visual and non-visual opsins in green anoles (*Anolis carolinensis*), and opsins were identified in the retina, pineal gland, and parietal eye [19].

Green anole lizards are among the most well-studied reptilian species and are thus ideally suited for investigations of non-visual opsins. Green anoles are routinely used in laboratory and field studies of morphology, ecology, behavior, and physiology (reviewed in [27, 26, 40]), and this species was the first reptile to have its genome sequenced and annotated [1, 12]. Green anoles are diurnal, arboreal lizards that naturally occur in the southeastern United States, with an invasive range that includes Guam and Hawaii, as well as Japan [42]. The anole breeding season (approximately late March to late July) is defined by photoperiod and temperature [23, 24], and this species primarily communicates using visual signals [17]. Nuanced detection of varying wavelengths of light, via visual and non-visual opsins, is likely to be essential for anole lizards.

## Results

Following PCR amplification, expression of OPN3-1, OPN3-3, OPN3-4, OPN4-1, OPN4-2, OPN5-2, and OPN5-6 was confirmed (see Figure 1a). Vertebrate ancient opsin (VAO) expression was also confirmed, but its expression was weaker as depicted by the faint band on the agarose gel. The rest of the transcripts could not be confirmed by RT-PCR amplification despite multiple attempts to optimize PCR reaction conditions and primer design. GAPDH was used as a control to show cDNA integrity and qPCR results showed dynamic expression of the confirmed transcripts. Confirmed opsins were most abundantly expressed in testes and liver, followed by eye and brain tissues, whilst skin showed lower expression levels. Eye tissue showed low values of all opsins except for OPN3-1 and OPN4-1, whilst the brain had very low levels of VAO and OPN5-6. In skin tissues, the dorsal skin consistently exhibited greater expression of all transcripts of interest than ventral and lateral skin (Figure 1b and c).

**Figure 1.**
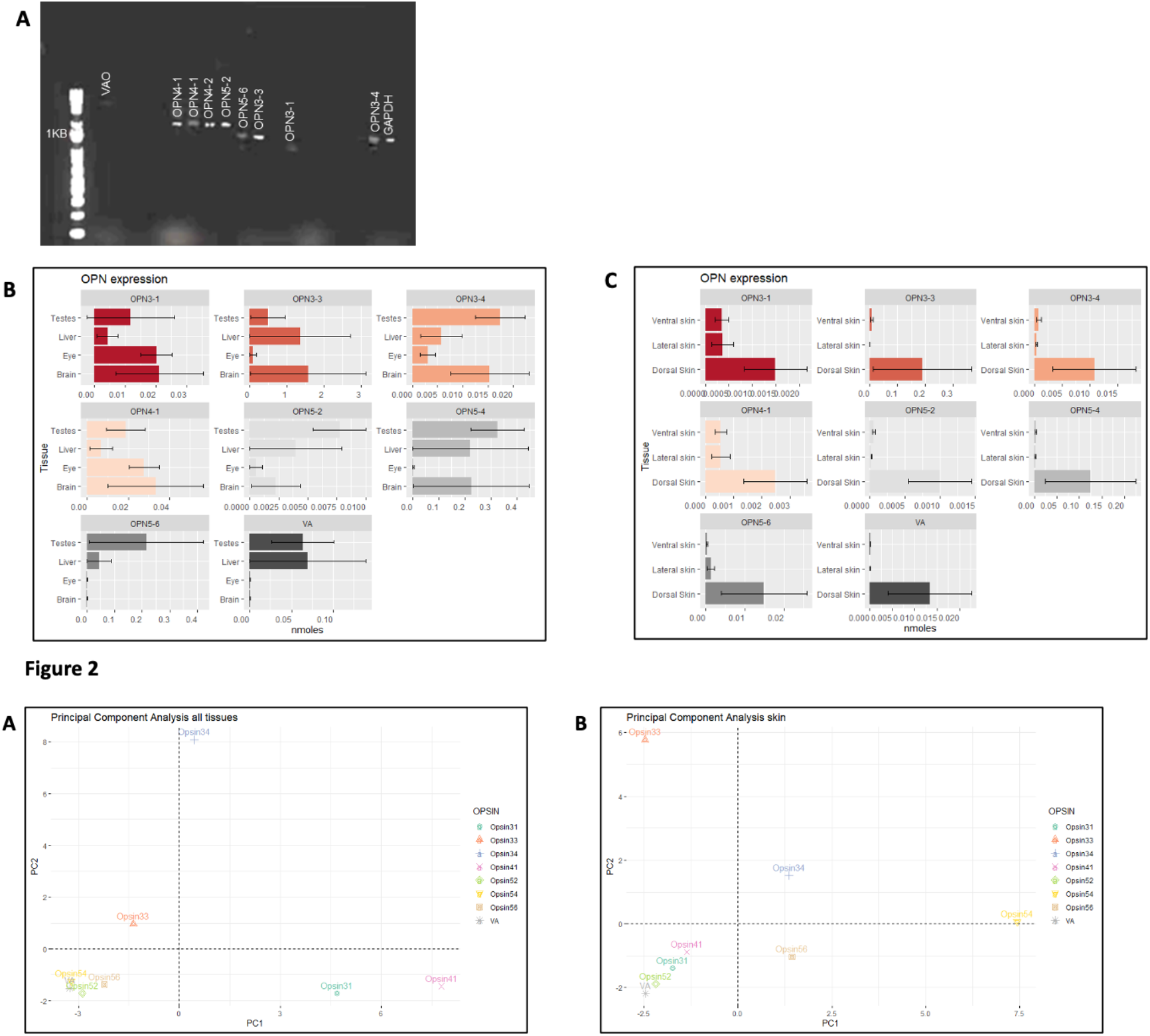
(a) Electrohopresis analysis of RT PCR targeting full green anole opsin transcripts. Bands show presence of confirmed transcripts. (b) qPCR results showing opsin expression levels in testes, liver, eye and brain tissues. (c) qPCR results of opsin expression in skin (ventral, lateral and dorsal) tissue.

### Differential expression among the opsins

OPN5-2, OPN3-4, OPN3-1 and OPN4-1 showed significant changes in expression levels among brain, testes, and eyes. The other opsin transcripts did not show any differences in expression among tissue types. Both OPN5 isoforms and OPN3-3 were the most expressed transcripts within brain, eyes, liver and testes. Within these transcripts, OPN5 was mostly expressed in testes whilst OPN3 expression was higher in the brain. PCA plots of opsin expression (Figure 2A) identified two clusters when comparing opsin expression in testes, liver, brain and eye (Figure 2A). The first cluster was composed of OPN5-4, OPN5-2, and OPN5-6 whilst OPN3-1 and OPN4-1 formed another cluster. OPN3-4 and OPN3-3 both showed expression patterns that were different to the rest of the opsins and their expression patterns did not cluster with any of the two groups described above. In the skin tissue a different pattern was observed (Figure 2B): OPN4-1, OPN3-1, OPN5-2 and VAO formed one cluster whilst OPN3-4, OPN5-6 and OPN5-4 showed slight similar expression. OPN3-3 did not show similar expression levels to any of the other opsins. Results are summarised in Table 2.0

**Figure 2.** (a) PCA plot of opsins based on qPCR expression in eyes, brain, testes and liver tissue. (b) PCA plot of opsins based on qPCR expression in skin tissue.

### Sequence conservation of confirmed opsins

To explore the sequence conservation of the confirmed opsins, a protein sequence alignment was performed. Looking at the domains each amino acid codes for, most of the 7 helixes conforming opsins show a conserved sequence. A lack of conservation was observed in the amino acids serving as “bridges” between helixes (Figure 3).

**Figure 3.**
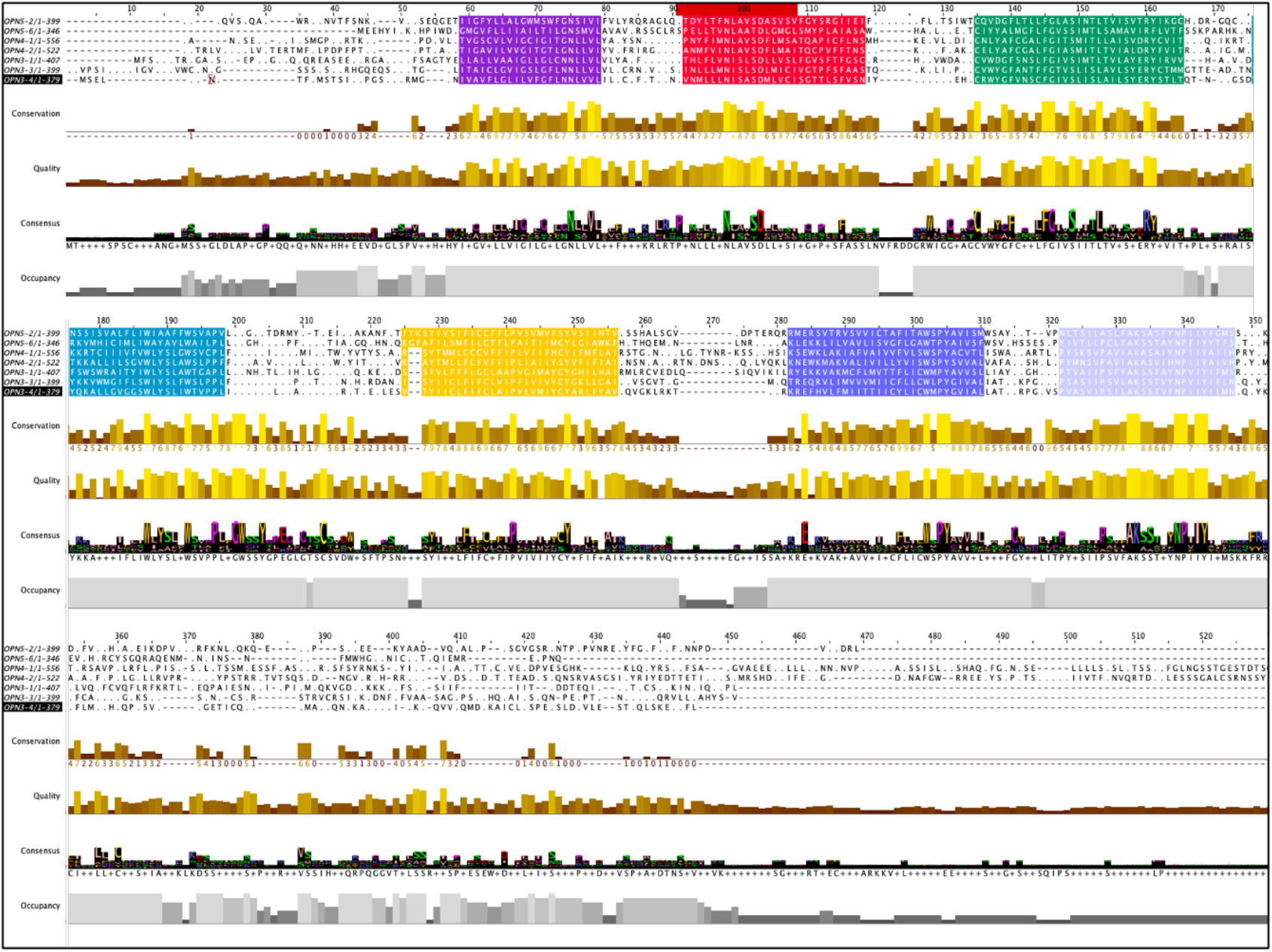
Protein alignment showing similarity of the green anole confirmed opsin transcripts. Each protein helix is depicted by a different color. Conservation levels are shown as bars underneath the alignment.

## Discussion

This study provides the first comprehensive evidence of non-visual opsin expression across the green anole body. Based on predicted sequences for different ERPs, non-visual opsin presence was confirmed in the brain, eye, testes, liver, and skin. We observed differences compared to other studies [34], with the highest level of expression of non-visual opsins observed in the liver and the testes, but not the brain, compared to the other tissues sampled.

### Encephalopsin (OPN3) family

High expression of ERPs, including OPN3, has been previously reported in the spermatids in the mammal testes [37]. OPN3 expression was also reported in the cerebral cortex and cerebellum, with lower levels in the eye, liver, and kidney [5]. These results are similar to the findings presented here for the green anole lizard. Other studies have reported important roles for OPN3 in the visual system of mice [25, 45], and our results corroborate a potential role for OPN3 in light reception in the anole eye. Our findings also suggest a role for all OPN3-related transcripts in the brain, but the possible role of each of the OPN3 transcripts cannot be identified because our data lacks protein or gene localization analysis. Interestingly, OPN3-3 was found to be the opsin with the highest level of expression in all of the tissues investigated, despite this not being the opsin with closer homology to mammal OPN3 (only 65% of the amino acids are conserved between mouse OPN3 and lizard OPN3-3). Future work will determine which specific brain structures express OPN3 to elucidate whether this gene has a conserved function between reptiles, birds, and mammals.

### Neuropsin (OPN5) family

This is the first study to report OPN5 expression in the testes of reptiles, and OPN5 was expressed at its highest levels in this tissue. Further studies will be necessary to determine the role that this opsin is playing in the reproductive system of the green anole. Previous studies have reported OPN5 ortholog expression in the mouse testes [45, 21], but the role of OPN5 in the reptilian testes is yet to be established. OPN5 expression has been previously identified in the hypothalamus of birds, fish, and mammals [35], but precise OPN5 localization in the lizard brain is yet to be reported. Although OPN5 paralogs have been previously identified, and confirmed by this study, OPN5-6 shows little expression in the brain suggesting a different role for this particular transcript.

### Melanopsin (OPN4) family

OPN4-1 was the only confirmed OPN4 family member that showed moderate expression in all green anole tissues. The most abundant expression of OPN4 was found in the brain and eye, suggesting a role in the light reception in the visual system as well as for deep-brain photoreception. Previous studies have reported OPN4 expression in the amphibian and bird hypothalamus, as well as in the premaxillary nucleolus of fish [39]. In lizards, an OPN4 ortholog has been successfully isolated from the telencephalon [34], but no other published studies have explored its expression in other brain regions. In mammals, OPN4 is expressed in the retinal ganglion cells of eyes, and it has been hypothesized that it acts as a light receptor that further transmits light cues to the brain [33]. The results from the current study support the hypothesis of OPN4 having a role in photodetection in the retina of anoles.

### Vertebrate ancient opsin (VAO)

VAO, contrary to what has been reported in other species, was expressed at low levels in the green anole brain. In birds, VAO has been reported to be expressed in the hypothalamus and has been shown to play an important role in the avian photoperiodic response [10, 21, 35, 36]. Previous studies in pinealectomized and retinalectomized ruin lizards (*Podarcis sicula*) have shown that inactivation of VAO, through administration of antisense mRNA, causes free-running locomotive activity, suggesting that photoperiodic entrainment of daily locomotor rhythms may be controlled by brain-based ERPs [13]. In the present study, VAO expression was not detected in the eyes, but it was expressed in the liver and testes.

### Differential expression of non-visual opsins in the skin

Opsins have been previously reported to be expressed in the skin of fish, amphibians, and reptiles, but one notable finding of this study is the consistent pattern of increased opsin expression in the dorsal skin compared to the lateral and ventral skin. Previous studies have reported the role of opsins in relation to skin color changes in fish (*Larimichthys crocea*) [51], and chameleons (species unidentified) [52]. Both species were able to maintain their color-changing abilities independently of visual inputs. OPN1 has also been detected in the tails of tadpoles (*Xenopus laevis*) [30] and the neon tetra (*Paracheirodon innesi*) [18], and more than 25 opsin variants have been found in the skin of zebrafish (*Danio rerio*) [20]. This study is the first to report a distinctive opsin expression pattern in the lizard skin.

Green anoles can change color from bright green to dark brown in response to stress, social cues, or thermoregulation [16]. This color change occurs in both the dorsal and lateral skin, but not in the ventral skin. This transformation is determined by the higher distribution of chromatophores that reflect light in the dorsal and lateral skin [46]. The findings from this study suggest that opsins may be present at higher concentrations where the chromatophores are present. Higher opsin expression in dorsal skin suggests that opsins are either mediating light responses in skin color changes or that light induces opsin expression so that higher opsin expression is found in skin receiving the highest light intensity. Our findings suggest that OPN3-3 is the most abundant opsin in the skin, and its expression levels also clustered separately from the other opsin families in the PCA plot. In mammals and birds, OPN3 has been identified as the opsin with the highest expression in skin [31]. This might suggest a conserved role in lizards.

To date, there have been no published studies reporting whether green anole lizards are capable of retaining their color-changing abilities in the absence of opsins. Opsins have also been related to sensory perception and dermal phototaxis. It has been reported that sea snakes (Hydrophiinae) have tails adapted to act as light reception organs by expressing photoreceptors such as RH1, SWS1, OPN5, and LWS [8]. It was suggested in this study that these snakes may have evolved this adaptation to allow photoreception in the tails when their heads are concealed, providing better environmental awareness. This is possibly similar to dermal photoreception observed in hogfish (*Lachnolaimus maximus*), where SWS1 opsin expression is found underneath chromatophores and subject to light changes from pigment activity [41]. However, there are no studies focusing on opsin expression in different skin types (dorsal, lateral, ventral). Our results did not detect a significant difference in opsin expression levels between lateral and ventral skin. This suggests that rather than light levels inducing opsin expression, there may be a specific group of cells expressing opsins in the dermis of green anoles.

### Opsins show similar sequences despite dynamic expression

Opsins are a class of transmembrane proteins that attach to the cell membrane through seven helix-type domains. These proteins are critical for photoreception and other light-sensitive processes. Although the structure of extra-retinal photoreceptors remains poorly understood, they exhibit substantial structural similarity to visual opsins, which have been extensively studied. Our study confirmed the sequence and potential structural conservation of opsins across species and tissue types. Analysis of green anole opsins revealed highly conserved sequences coding helix structures, despite these opsins being expressed in a wide range of tissues. This raises an intriguing question: how do opsins with such similar sequences [Supplementary Tables 1-3] and conserved structural features differentiate to perform specialized functions in distinct tissues? One possible explanation is that the specific roles of each opsin are defined by non-conserved amino acids within the helices or by unique “bridging” elements between helices that influence their activity and localization.

This study supports the idea that conserved opsin structures may underpin shared fundamental functions. For example, similar structural motifs in visual and non-visual opsins across vertebrates suggest that key aspects of their mechanisms are preserved. However, the observed variation in opsin expression patterns across tissues points to an additional layer of regulation, potentially driven by differences in tissue-specific cofactors, post-translational modifications, or interactions with unique cellular environments. In green anoles, the high sequence similarity and shared amino acid motifs among opsins and their isoforms make them a compelling model for studying these phenomena. Ultimately, while opsin structure is highly conserved, both within lizards and across other species, functional specialization is likely dictated by subtle differences in sequence and the microenvironment of each tissue. These findings set the basis for further investigations into the evolutionary and functional dynamics of opsins in diverse biological systems.

## Methods

### Identification of opsin sequences in the green anole genome

In an analysis of the evolution of opsin genes across vertebrates, Beaudry and colleagues [3] identified 13 opsin gene sequences in the *Anolis carolinensis* genome. To date, this published compilation provides the most comprehensive list of anole opsin sequences. In order to match these previously identified opsins [3] with the green anole annotations available at the time of this study, distinctions between visual and non-visual opsins were made. Ten transcripts were identified as non-visual opsins [3]; the remaining three were recognized as visual opsins. Following identification, a BLASTN was conducted against the gene sequences of the non-visual opsins on both the ENSEMBL and the NCBI databases using the green anole AnoCar2.0v2 (genomic sequence) as a reference. When this study was conducted in November 2022, ENSEMBL recognized only one sequence in each of the opsin 3, 4, and 5 gene families (Table 1), but did not identify the other seven sequences as opsin gene sequences. In contrast, the NCBI blast returned predicted sequences for all the opsins except the vertebrate ancient opsin (VAO), which had been validated by RNA-seq. Isoforms for several non-visual opsin genes were also predicted in NCBI (Table 1), but were not consistently identified in ENSEMBL. Both databases confirmed that all sequences predicted to be opsins are orthologous to sequences identified as opsins in other taxa. For example, ENSEMBL showed that these sequences have 1:1 orthologues within Sauropsida (birds and reptiles). The predicted transcripts on both ENSEMBL and NCBI were summarized and primers designed to amplify them.

**Table 1.**
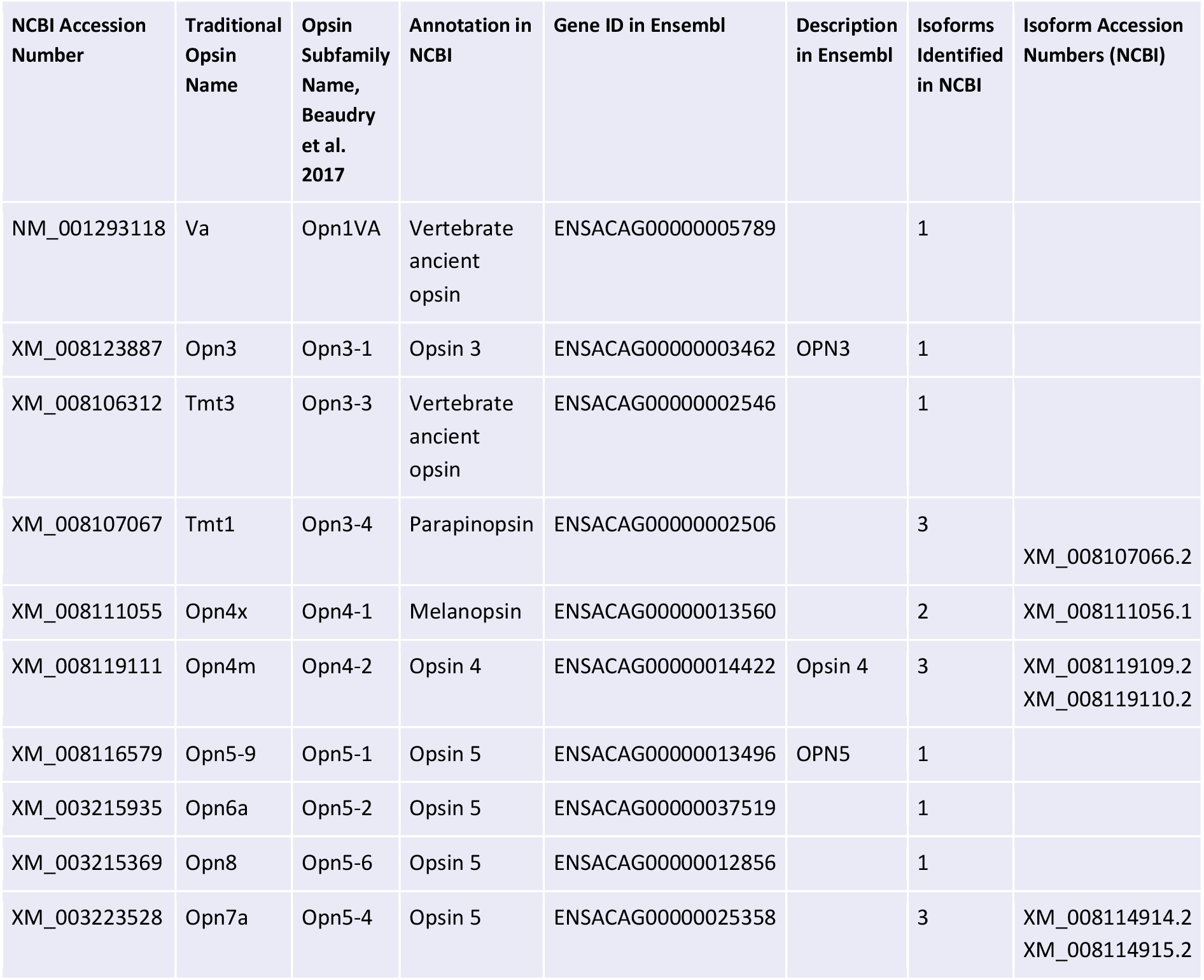
Predicted green anole (*Anolis carolinensis*) opsin sequences found on ENSEMBL and/or NCBI.

**Table 2:**
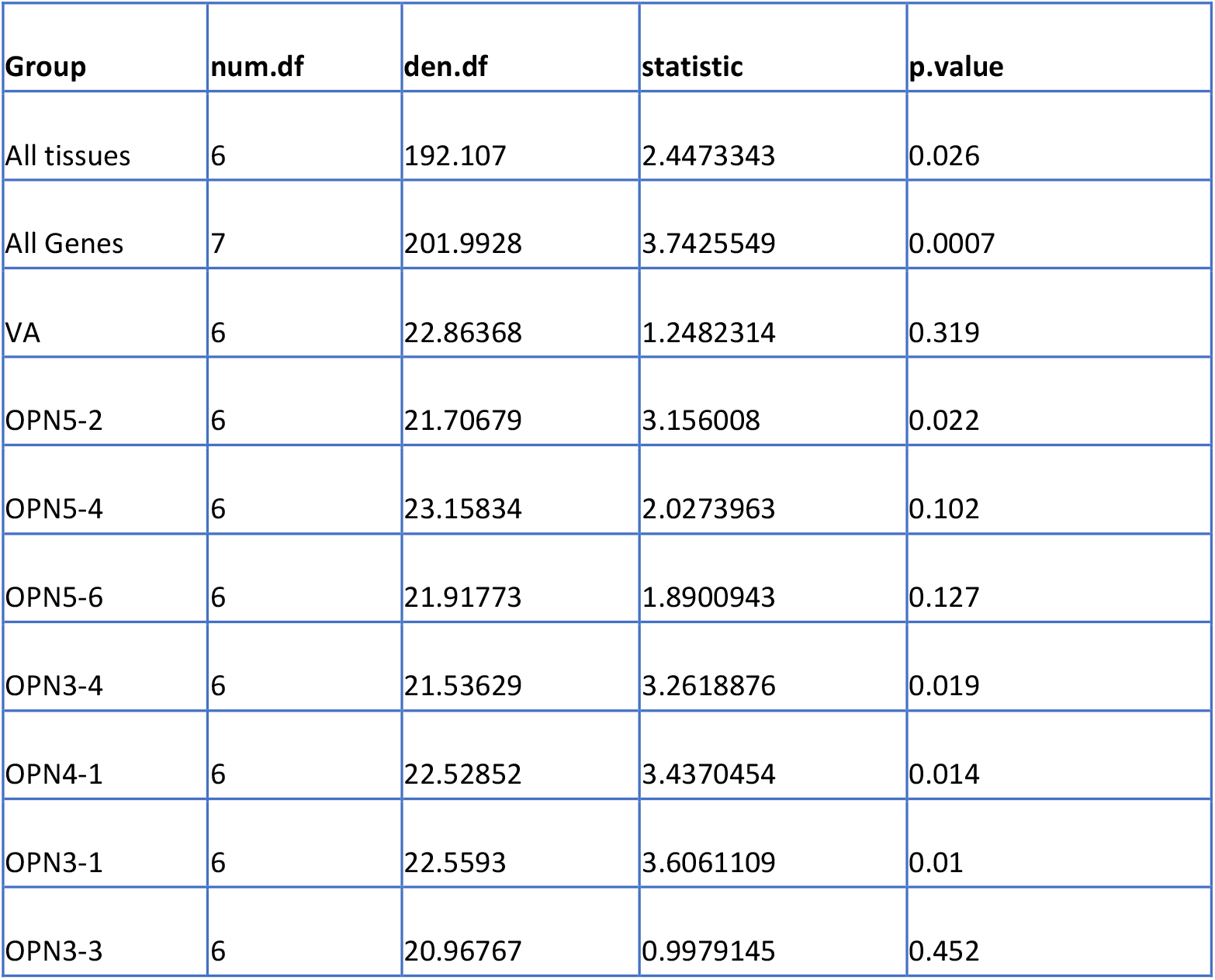
Statistical results of OPNX expression comparison between different tissues of the green anole (*Anolis carolinensis*).

### Alignment of opsin sequences and phylogeny

Within each of the three non-visual opsin subfamilies (OPN3, OPN4, and OPN5), a multiple alignment of the coding sequences was performed using the default parameters in CLUSTALW in MEGA X [22]. These alignments allowed for the visualization of similarities and differences among predicted isoforms within a gene and across paralogous sequences within an opsin subfamily. A gene tree of all green anole opsin coding sequences (10 non-visual opsins, 3 visual opsins, and their predicted isoforms) was constructed for a total of 1934 nucleotide positions using Maximum Likelihood and the Tamura-Nei model [43] with 500 bootstrap replicates (Supplementary Figure 1). The alignment in CLUSTALW at the amino acid level was repeated, and the results visualized in Jalview 2.11.3.2. Protein domain search was performed using the NCBI conserved domain search tool (https://www.ncbi.nlm.nih.gov/Structure/cdd/wrpsb.cgi).

### Primer design and sequencing validation

Primers were designed based on the sequences found on the NCBI genome browser (https://www.ncbi.nlm.nih.gov/), and oligonucleotides were made to target the various paralog genes. To confirm the NCBI sequences, primers were designed to amplify whole transcripts to target 200–300 bp of the validated transcripts to quantify their expression by qPCR. Primer pairs targeting each of the paralogs were designed to avoid isoform-specific sequences and isoforms (see Supplementary Table 4). Glyceraldehyde-3-phosphate dehydrogenase (GAPDH) was used as a housekeeping gene. Primer pairs were first tested by PCR to ensure they amplified a single product, and amplicon sequences of the products were confirmed by The University of Edinburgh Sequencing Facility (UK). For each primer pair, a full-transcript amplification was attempted, and the primer sequence and/or the PCR reaction was optimized if it was unsuccessful. For some transcripts, amplification was not achieved despite smaller amplicon amplification, suggesting that these transcripts were either not expressed in the selected tissues, the predicted sequence was not correct, or they are not real transcripts (i.e., the prediction is incorrect). All agarose gels were visualised in a iBright CL750 Imaging System and annotated using ImageJ. Constrat was adjusted to be able to visualise all existing bands.

### Tissue collection and preparation of cDNA

Adult green anole lizards (4 females and 6 males) were captured during daylight hours on the campus of Trinity University in San Antonio, Texas, USA in mid-April 2023, approximately at the beginning of the breeding season for this species. Green anole lizard collection was performed under Scientific Permit SPR-0310-045 to Michele A. Johnson from Texas Parks and Wildlife Department, with approval from Trinity University’s Animal Research Committee, protocol 051122-MJ2. Each lizard was caught using a dental floss loop and placed in an individual cloth bag for transport to the laboratory, where it was euthanized using the two-stage MS-222 protocol (intracoelomic injection of 250–500 mg/kg pH-neutralized 1% MS-222, followed by 0.5–1 ml 50% unbuffered MS-222; [7]). Following euthanasia, brain, eyes, liver, and gonadal tissues were removed. During dissections, the pineal gland was not removed from the brain. Skin samples (∼25 mm^2^) were also collected from the middle third of the ventral region (belly), left and right lateral regions, and the dorsum (back) of each lizard. All tissues were immediately transferred to 1.5 ml tubes containing 1 ml of RNAlater (Sigma Aldrich, USA) and maintained on a shaker at 4°C overnight, then stored at 4°C for 48 h. Tissue samples were then shipped overnight, at ambient temperature, to The Roslin Institute, The University of Edinburgh, UK.

RNA was extracted from each tissue sample (brain, eye, testis, ovary, liver, dorsal skin, lateral skin, and ventral skin) using a RNeasy Mini Kit (Qiagen, Germany), following the manufacturer’s instructions. Tissue was homogenized using a handheld homogenizer. RNA was eluted in 30 μl RNase-free water, and quality and concentration were confirmed by running each sample on a 1% agarose gel and Nanodrop. Total RNA was stored at −80°C and the samples were then diluted to 300 ng/ml to synthesize cDNA. A high-capacity cDNA reverse transcription kit (ThermoFisher, USA) was used to synthesize cDNA using random primers, without RNase inhibitor, following the manufacturer’s instructions; cDNA was stored at 4°C until use.

### Transcript expression

RT-PCR was carried out using Phire Hot Start II DNA Polymerase (ThermoFisher, USA) following the manufacturer’s instructions to confirm the transcripts predicted by NCBI; 500 ng of pooled (all tissues) cDNA was used as starting material. The thermal profile for this reaction consisted of initial denaturation at 98°C for 30 s, followed by 35 cycles of denaturation at 98°C for 5 s and annealing at 63.5°C for 15 s, and a final extension period at 72°C for 1 min. Following PCR, all products were run on a 2% agarose gel to confirm expression and check for alternative isoforms. Transcripts that were confirmed by RT-PCR were then selected to measure their expression in selected tissues. Primers amplifying 200–300 bp amplicons were first tested by RT-PCR and products run on a 1% agarose gel. Products were then cleaned using a PCR purification kit (QIAGEN Ltd., UK). The concentration was measured using Nanodrop. Standard curves for each primer pair were prepared by diluting the PCR-cleaned product by 1:10 (8 points per standard curve). Once standard curves were constructed, qPCR reactions on samples of interest were carried out using qRT-PCR Brilliant III SYBR Master Mix (Agilent Technologies, Cedar Creek, Texas, USA) on an MX3000Pro, following the manufacturer’s instructions. ROX (Agilent Technologies, Cedar Creek, Texas, USA) was used as a reference dye, and all reactions were analyzed using MXPro software (Agilent Technologies, Cedar Creek, Texas, USA) by calculating the total number of moles in each sample using the standard curve as a reference. All values were normalized using GAPDH as a reference housekeeping gene.

### Protein sequence alignment

Protein alignments were performed by retrieving the protein sequences from the NCBI protein database (https://www.ncbi.nlm.nih.gov/protein/) and aligning them using Clustal Omega Multiple Sequence Aligner (https://www.ebi.ac.uk/Tools/msa/clustalo/). Results were visualized using Jalview [50].

### Statistical analysis

Statistical analysis was performed using R 4.2.2 (https://www.r-project.org/) with the package *stats*. Biological and technical replicates were used (i.e., a total of five samples from individual lizards were run in triplicate), regardless of the sex, to calculate the geometric mean. To test for differences in expression at three levels, a one-way ANOVA was carried out for the following comparisons:

1. Differences within all tissues (i.e., opsin expression differences in dorsal vs. ventral skin, brain vs. liver, etc.).
2. Among all genes (i.e., expression levels of OPN5-1 in brain, liver, eyes, etc., vs. OPN5-2 expression levels in brain, liver, eyes, etc.).
3. Individual gene expression in all tissues (i.e., expression of OPN5-1 in brain compared to eyes, liver, etc.).

Results were defined as significant if the p-value was ≤ 0.05. To explore expression patterns of the opsins, a PCA analysis approach in R was used to dissect the similarity levels between them, after normalizing the data with the *scale* function and subsequently applying the *princomp* function. The data were visualized to explore the clusters formed by the genes of interest. These clusters confirmed which opsins had similar expression patterns in each tissue.

## Data availability

The datasets used during the current study are available at https://zenodo.org/records/14744958

## Acknowledgements

This work was supported by an International Institutional Award to the University of Edinburgh (BB/Y51410X/1) and Roslin Institute Strategic Grant (BBS/E/RL/230001C) funding from the UK Biotechnology and Biological Sciences Research Council to Simone L. Meddle along with financial support from the Trinity University Office of Academic Affairs to Michele A. Johnson. We would like to thank Dale Cochran and members of the Johnson Lab for all of their fantastic help in the laboratory and field.

## Author contributions

V.T-R, M.A.J, and S.L.M. designed the research; V.T-R, M.A.J, G.E.A and A.R.J. performed the research and V.T-R, M.A.J, and A.R.J. analyzed the data; V.T-R, M.A.J, and S.L.M. wrote the manuscript. All the authors were involved in drafting and revising the manuscript.

## References

1. Alföldi, J., Di Palma, F. & Grabherr, M. The genome of the green anole lizard and a comparative analysis with birds and mammals. Nature 477, 587–591. 10.1038/nature10390 (2011).

2. Bailey, M. J. & Cassone, V. M. Opsin photoisomerases in the chick retina and pineal gland: characterization, localization, and circadian regulation. Invest. Ophthalmol. Vis. Sci. 45, 769–775. 10.1167/iovs.03-1125 (2004).

3. Beaudry, F. E. G. et al. The non-visual opsins: eighteen in the ancestor of vertebrates, astonishing increase in ray-finned fish, and loss in amniotes. J. Exp. Zool. Part B Mol. Dev. Evol. 328, 685–696. 10.1002/jez.b.22764 (2017).

4. Bertolesi, G. E., Debnath, N., Malik, H. R., Man, L. L. H. & McFarlane, S. Type II opsins in the eye, the pineal complex and the skin of Xenopus laevis: using changes in skin pigmentation as a readout of visual and circadian activity. Front. Neuroanat. 15, 784478. 10.3389/fnana.2021.784478 (2021).

5. Blackshaw, S. & Snyder, S. H. Encephalopsin: a novel mammalian extraretinal opsin discretely localized in the brain. J. Neurosci. 19, 3681–3690 (1999).

6. Chong, Y. H., Kitahashi, T. & Parhar, I. S. Neuronal organization of deep brain opsin photoreceptors in adult teleosts. Front. Neuroanat. 10, 48. 10.3389/fnana.2016.00048 (2016).

7. Conroy, C. J., Papenfuss, T., Parker, J. & Hahn, N. Use of tricaine methanesulfonate (MS-222) for euthanasia of reptiles. J. Am. Assoc. Lab. Anim. Sci. 48, 28–32 (2009).

8. Crowe-Riddell, J. M. et al. Phototactic tails: evolution and molecular basis of a novel sensory trait in sea snakes. Mol. Ecol. 28, 2013–2028. 10.1111/mec.15022 (2019).

9. Davies, W. I. L. et al. An extended family of novel vertebrate photopigments is widely expressed and displays a diversity of function. Genome Res. 25, 1666–1679. 10.1101/gr.189886.115 (2015).

10. Davies, W. I. L. et al. Vertebrate ancient opsin photopigment spectra and the avian photoperiodic response. Biol. Lett. 8, 445–448. 10.1098/rsbl.2011.0864 (2012).

11. Dong, C. M., McLean, C. A., Moussalli, A. & Stuart-Fox, D. Conserved visual sensitivities across divergent lizard lineages that differ in an ultraviolet sexual signal. Ecol. Evol. 9, 11824–11832. 10.1002/ece3.5686 (2019).

12. Eckalbar, W. L. et al. Genome reannotation of the lizard Anolis carolinensis based on 14 adult and embryonic deep transcriptomes. BMC Genomics 14, 49. 10.1186/1471-2164-14-49 (2013).

13. Foà, A., Flamini, M., Innocenti, A., Minutini, L. & Monteforti, G. The role of extraretinal photoreception in the circadian system of the ruin lizard Podarcis sicula. Comp. Biochem. Physiol. A Physiol. 105, 223–230. 10.1016/0300-9629(93)90199-E (1993).

14. Fulgione, D. et al. Seeing through the skin: dermal light sensitivity provides cryptism in Moorish gecko. J. Zool. 294, 122–128. 10.1111/jzo.12155 (2014).

15. García-Fernández, J. M. et al. The hypothalamic photoreceptors regulating seasonal reproduction in birds: a prime role for VA opsin. Front. Neuroendocrinol. 37, 13–28. 10.1016/j.yfrne.2014.11.001 (2015).

16. Horr, D. M., Payne, A. A., McEntire, K. D. & Johnson, M. A. Sexual dimorphism in dynamic body color in the green anole lizard. Behav. Ecol. Sociobiol. 77, 34. 10.1007/s00265-023-03286-3 (2023).

17. Jenssen, T. A. Evolution of anoline display behavior. Am. Zool. 17, 203–215 (1977).

18. Kasai, A. & Oshima, N. Light-sensitive motile iridophores and visual pigments in the neon tetra, Paracheirodon innesi. Zoolog. Sci. 23, 815–819. 10.2108/zsj.23.815 (2006).

19. Kawamura, S. & Yokoyama, S. Expression of visual and nonvisual opsins in American chameleon. Vision Res. 37, 1867–1871. 10.1016/S0042-6989(96)00309-4 (1996).

20. Kelley, J. L. & Davies, W. I. The biological mechanisms and behavioral functions of opsin-based light detection by the skin. Front. Ecol. Evol. 4, 106. 10.3389/fevo.2016.00106 (2016).

21. Kojima, D. et al. UV-sensitive photoreceptor protein OPN5 in humans and mice. PLoS ONE 6, e26388. 10.1371/journal.pone.0026388 (2011).

22. Kumar, S., Stecher, G., Li, M., Knyaz, C. & Tamura, K. MEGA X: molecular evolutionary genetics analysis across computing platforms. Mol. Biol. Evol. 35, 1547–1549. 10.1093/molbev/msy096 (2018).

23. Licht, P. Regulation of the annual testis cycle by photoperiod and temperature in the lizard Anolis carolinensis. Ecology 52, 240–252. 10.2307/1934178 (1971).

24. Licht, P. Influence of temperature and photoperiod on the annual ovarian cycle in the lizard Anolis carolinensis. Copeia 1973, 465–472. 10.2307/1443116 (1973).

25. Linne, C. et al. Encephalopsin (OPN3) is required for normal refractive development and the GO/GROW response to induced myopia. Mol. Vis. 29, 39–57. PMID: 37287644; PMCID: PMC10243678 (2023).

26. Losos, J. B. Lizards in an evolutionary tree: ecology and adaptive radiation of anoles. Univ. of California Press (2011).

27. Lovern, M. B., Holmes, M. M. & Wade, J. The green anole (Anolis carolinensis): a reptilian model for laboratory studies of reproductive morphology and behavior. ILAR J. 45, 54–64. 10.1093/ilar.45.1.54 (2004).

28. Mano, H., Kojima, D. & Fukada, Y. Exo-rhodopsin: a novel rhodopsin expressed in the zebrafish pineal gland. Mol. Brain Res. 73, 110–118. 10.1016/S0169-328X(99)00174-5 (1999).

29. Miyashita, Y. et al. The photoreceptor molecules in Xenopus tadpole tail fin, in which melanophores exist. Zoolog. Sci. 18, 671–674. 10.2108/zsj.18.671 (2001).

30. Miyashita, Y., Moriya, T., Kubota, T., Yamada, K. & Asami, K. Expression of opsin molecule in cultured murine melanocyte. J. Investig. Dermatol. Symp. Proc. 6, 54–57. 10.1046/j.0022-202x.2001.00018.x (2001).

31. Olinski, L. E., Tsuda, A. C. & Oancea, E. Endogenous opsin 3 (OPN3) protein expression in the adult brain using a novel OPN3-mCherry knock-in mouse model. eNeuro 7, ENEURO.0107-20.2020. 10.1523/ENEURO.0107-20.2020 (2020).

32. Palczewski, K. & Kiser, P. D. Shedding new light on the generation of the visual chromophore. Proc. Natl Acad. Sci. USA 117, 19629–19638. 10.1073/pnas.2008211117 (2020).

33. Pan, J. Q. et al. Long-day photoperiods affect expression of OPN5 and the TSH-DIO2/DIO3 pathway in Magang goose ganders. Poult. Sci. 101, 102024. 10.1016/j.psj.2022.102024 (2022).

34. Pasqualetti, M. et al. Identification of circadian brain photoreceptors mediating photic entrainment of behavioural rhythms in lizards. Eur. J. Neurosci. 18, 364–372. 10.1046/j.1460-9568.2003.02770.x (2003).

35. Pérez, J. H. et al. A comparative perspective on extra-retinal photoreception. Trends Endocrinol. Metab. 30, 39–53. 10.1016/j.tem.2018.10.001 (2019).

36. Pérez, J. H. et al. Functional inhibition of deep brain non-visual opsins facilitates acute long day induction of reproductive recrudescence in male Japanese quail. Horm. Behav. 145, 105298. 10.1016/j.yhbeh.2022.105298 (2023).

37. Pérez-Cerezales, S. et al. Involvement of opsins in mammalian sperm thermotaxis. Sci. Rep. 5, 16146. 10.1038/srep16146 (2015).

38. R Core Team. R: a language and environment for statistical computing. R Foundation for Statistical Computing. https://www.R-project.org (2022).

39. Salgado, D. et al. Light-induced shifts in opsin gene expression in the four-eyed fish Anableps anableps. Front. Neurosci. 16, 995469. 10.3389/fnins.2022.995469 (2022).

40. Sanger, T. J. & Kircher, B. K. Model clades versus model species: Anolis lizards as an integrative model of anatomical evolution. In: Sheng, G. (Ed.) Avian and Reptilian Developmental Biology. Methods in Molecular Biology 1650, 1–19. 10.1007/978-1-4939-7216-6_1 (2017).

41. Schweikert, L. E. et al. Dynamic light filtering over dermal opsin as a sensory feedback system in fish color change. Nat. Commun. 14, 4642. 10.1038/s41467-023-40166-4 (2023).

42. Suzuki-Ohno, Y. et al. Factors restricting the range expansion of the invasive green anole Anolis carolinensis on Okinawa Island, Japan. Ecol. Evol. 7, 4357–4366. 10.1002/ece3.2992 (2017).

43. Tamura, K. & Nei, M. Estimation of the number of nucleotide substitutions in the control region of mitochondrial DNA in humans and chimpanzees. Mol. Biol. Evol. 10, 512–526. 10.1093/oxfordjournals.molbev.a040023 (1993).

44. Taniguchi, Y., Hisatomi, O., Yoshida, M. & Tokunaga, F. Evolution of visual pigments in geckos. FEBS Lett. 445, 36–40. 10.1016/S0014-5793(99)00089-7 (1999).

45. Tarttelin, E. E., Bellingham, J., Hankins, M. W., Foster, R. G. & Lucas, R. J. Neuropsin (Opn5): a novel opsin identified in mammalian neural tissue. FEBS Lett. 554, 410–416. 10.1016/S0014-5793(03)01265-4 (2003).

46. Taylor, J. D. & Hadley, M. E. Chromatophores and color change in the lizard, Anolis carolinensis. Z. Zellforsch. Mikrosk. Anat. 104, 282–294. 10.1007/BF00309737 (1970).

47. Terakita, A. The opsins. Genome Biol. 6, 213. 10.1186/gb-2005-6-3-213 (2005).

48. Tseng, W. H. et al. Opsin gene expression regulated by testosterone level in a sexually dimorphic lizard. Sci. Rep. 8, 16055. 10.1038/s41598-018-34284-z (2018).

49. Underwood, H. & Menaker, M. Extraretinal light perception: entrainment of the biological clock controlling lizard locomotor activity. Science 170, 190–193. 10.1126/science.170.3953.190 (1976).

50. Waterhouse AM, Procter JB, Martin DMA, Clamp M, Barton GJ Jalview Version 2 - A multiple sequence alignment editor and analysis workbench. Bioinformatics 25 1189–1191. (doi:10.1093/bioinformatics/btp033) (2009)

51. Zhang, Z. et al. Nonvisual system-mediated body color change in fish reveals nonvisual function of Opsin 3 in skin. J. Photochem. Photobiol. B. 252, 112861. 10.1016/j.jphotobiol.2024.112861 (2024).

52. Zoond, A. & Bokenham, N. Studies in reptilian colour response: the role of retinal and dermal photoreceptors in the pigmentary activity of the chameleon. J. Exp. Biol. 12, 39–43 (1935).

