## Supplementary tables for "Expression of Non-Visual Opsins in the Green Anole Lizard (*Anolis carolinensis*)"

### Supplementary Information

**Trejo-Reveles et al.**

**Supplementary Table 1.**

Alignment of *Anolis carolinensis* coding sequences in the Opsin 3 gene family, including three paralogs (OPN3-1, OPN3-3, and OPN3-4), and three predicted isoforms of OPN3-4.

```
#NEXUS
[ Title Phylogenetic Analysis, Opsin 3]
begin data;
   dimensions ntax=5 nchar=1245;
   format    missing=?    gap=-    matchchar=.    datatype=nucleotide
interleave=yes;
   matrix

[!Domain=Data property=Coding CodonStart=1;]
OPN3_1  ATGTTCTCCGCCAACGGGACCCGGAGCGGCGCCGGCAGCGAC-CTGGAGCCGGGCCCGGG
OPN3_3  ...AC.GTGC.T.G.ATC.G..CC..TT..ATT..TGT..CGAAC..T.T.T..TG.A.-
OPN3_4A ----------------------------------------.TGTCA..A.TTA..T.---
OPN3_4B ----------------------------------------.TGTCA..A.TTA..T.---
OPN3_4C ----------------------------------------.TGTCA..A.TTA..T.---

OPN3_1  GCAGCAGCAGCAGCAGCGGGAGGCGTCGGAGGAGGAGGAGAGGGGAGCGGGGCTCTCGCC
OPN3_3  -..A.G..G.......TA.C.ACA.C.ATCACC.ACATG.GCAA....A.A..CTGA.T.
OPN3_4A -..ATTTG.CTTT..ATAT...CA-C.TC.ATT.A..AGCCA..A..T..ACTCTCAAGA
OPN3_4B -..ATTTG.CTTT..ATAT...CA-C.TC.ATT.A..AGCCA..A..T..ACTCTCAAGA
OPN3_4C -..ATTTG.CTTT..ATAT...CA-C.TC.ATT.A..AGCCA..A..T..ACTCTCAAGA

OPN3_1  CTTCAGCGCCGGGACCTACGAGCTGCTGGCCCTGCTGGTGGCCGCCATCGGCCTGCTGGG
OPN3_3  .CA.C.-...ACCT.A.CAC...CATCT...TC.G..-------T...T..GAGC.....
OPN3_4A --ATG.-.A.ACA..A.TGTC....T.TTT.T..GAC-------TA..TCTAG.TT.T..
OPN3_4B --ATG.-.A.ACA..A.TGTC....T.TTT.T..GAC-------TA..TCTAG.TT.T..
OPN3_4C --ATG.-.A.ACA..A.TGTC....T.TTT.T..GAC-------TA..TCTAG.TT.T..
```

```
OPN3_1   CTTGTGCAACAACCTGCTGGTGCTGGTGCTCTACGCCAAGTTCAAGCGGCTCCGGACGCC
OPN3_3   T..CCTG.......C.....T.....C..T.T.TG.CG.AAT..AGTT......T.T..
OPN3_4A G..C.TG..T...T..G.T..C..CA.T....T.TGT.....T..AACCT.G..C.AC..
OPN3_4B G..C.TG..T...T..G.T..C..CA.T....T.TGT.....T..AACCT.G..C.AC..
OPN3_4C G..C.TG..T...T..G.T..C..CA.T....T.TGT.....T..AACCT.G..C.AC..

OPN3_1   GACGCACCTCTTCCTGGTCAACATCAGCCTCAGCGACCTGCTGGTCTCCCTCTTCGGGGT
OPN3_3   A.TCA....GC.G...A.G.....TTCAT.G..T......A..A.A.G.A.TG.G..CAC
OPN3_4A TGT.A..A.GC.GT..C.G..T.....TGC...T...A....A....G.A...CT..AAC
OPN3_4B TGT.A..A.GC.GT..C.G..T.....TGC...T...A....A....G.A...CT..AAC
OPN3_4C TGT.A..A.GC.GT..C.G..T.....TGC...T...A....A....G.A...CT..AAC

OPN3_1   CAGCTTCACCTTCGGCTCCTGCCTCCGGCACCGCTGGGTCTGGGACGCCGCCGGGTGCGT
OPN3_3   .CCT....G.....CTG..A..ACG.A.GG.AAG...C..ATT.GGC.A..A..C..T..
OPN3_4A T.CAC...G.....TA..AAA.A.TTATGG.A.A...A..G.A.GA.AACA...C...AG
OPN3_4B T.CAC...G.....TA..AAA.A.TTATGG.A.A...A..G.A.GA.AACA...C...AG
OPN3_4C T.CAC...G.....TA..AAA.A.TTATGG.A.A...A..G.A.GA.AACA...C...AG

OPN3_1   CTGGGACGGCTTCAGCAACAGCCTTTTCGGAATTGTTTCTATCATGACACTCACCGTTCT
OPN3_3   ....T......TGC.....C.T.C..T..T.C...C..AC.G..AT.T..GG.T..G..
OPN3_4A G...T.......TGT...TTCTTGC..T..C.....A...C.G..CT....GG..A....
OPN3_4B G...T.......TGT...TTCTTGC..T..C.....A...C.G..CT....GG..A....
OPN3_4C G...T.......TGT...TTCTTGC..T..C.....A...C.G..CT....GG..A....

OPN3_1   GGCCTACGAACGTTACATCAGAGTTGTGCATGCAA---GAGTG------ATTGATTTCTC
OPN3_3   TT.A..T.....A...TG..C.A.GA..GGAA...CAGA..CAGATGCC.CAA...A.AA
OPN3_4A .T.T..T........T.G..CTC..ACA..AA.C.-ATA.AC.--TGGCTC...C.ATCA
OPN3_4B .T.T..T........T.G..CTC..ACA..AA.C.-ATA.AC.--TGGCTC...C.ATCA
OPN3_4C .T.T..T........T.G..CTC..ACA..AA.C.-ATA.AC.--TGGCTC...C.ATCA

OPN3_1   ATGGTCTTGGCGGGCAATCACATACATCTGGCTCTATTCCTTAGCCTGGACAGGGGCACC
OPN3_3   GAAAGTA...AT..GC...TTC.TGTCA...A.A.....T...TTT...T..CTAC....
OPN3_4A .AAAG.CCT..TA.GCG.TGG.GG.TC.........C..TC.GATT.......TAC.G..
```

```
OPN3_4B .AAAG.CCT..TA.GCG.TGG.GG.TC.........C..TC.GATT.......TAC.G..
OPN3_4C .AAAG.CCT..TA.GCG.TGG.GG.TC.........C..TC.GATT.......TAC.G..

OPN3_1  TCTCCTTGGCTGGAACCATTACACACTAGAAATACATGGATTAGGTTGTTCGGTGGACTG
OPN3_3  A..TT....A....GTAGC..TGG..CT...GG..CA..CACCACA.....T...A....
OPN3_4A ....A.........G.AGC..TGGG..T..GGGGGCG..CACCA.......A..CCGA..
OPN3_4B ....A.........G.AGC..TGGG..T..GGGGGCG..CACCA.......A..CCGA..
OPN3_4C ....A.........G.AGC..TGGG..T..GGGGGCG..CACCA.......A..CCGA..

OPN3_1  G-CAGTCGAAAGAGCCCAGCGATTCCTCCTTTGTTCTATTTTTCTTTCTTGGTTGTCTAG
OPN3_3  .-..T....GG..TG.T.A.A.CATA..T.ACA.CA.T.GCC.T..CA.CTT...C..T.
OPN3_4A .A..TCG....C.CT.G.ATC.-GTGA.T.A.A.CA.C.GCC....CA.ATTC..CT...
OPN3_4B .A..TCG....C.CT.G.ATC.-GTGA.T.A.A.CA.C.GCC....CA.ATTC..CT...
OPN3_4C .A..TCG....C.CT.G.ATC.-GTGA.T.A.A.CA.C.GCC....CA.ATTC..CT...

OPN3_1  CAGCCCCAGTTGGAATCATGGCTTACTGCTATGGGCATATTCTTCATGCAATACGAATGC
OPN3_3  TCATT..CT..ATCG.T..T.T...T........AA.GC.G..GTG...C..--------
OPN3_4A .CATT..C..CCTGG....ATA...........CT.GCC.CT..T....TG.--------
OPN3_4B .CATT..C..CCTGG....ATA...........CT.GCC.CT..T....TG.--------
OPN3_4C .CATT..C..CCTGG....ATA...........CT.GCC.CT..T....TG.--------

OPN3_1  TTCGATGTGTTGAAGACTTGCAATCTATCCAAGTGATCAAGATCCTAAGATATGAGAAGA
OPN3_3  --.A.GAA.G...GTG-----G.GTA.C.....GA..---.GCA.A..CGCGA...C.AC
OPN3_4A --.A.ACA.G..GG..-----..CTCCGAA..ACCTC---TGC..GG.A.AGA..ATTCC
OPN3_4B --.A.ACA.G..GG..-----..CTCCGAA..ACCTC---TGC..GG.A.AGA..ATTCC
OPN3_4C --.A.ACA.G..GG..-----..CTCCGAA..ACCTC---TGC..GG.A.AGA..ATTCC

OPN3_1  AAGTGGCTAAAATGTGTTTTTTAATGGTCACAACTTTTCTAATTTGTTGGATGCCTTATG
OPN3_3  G...CCTG.TC...GTGG..G.G...A...TTTGC.....GC....C...C.A.......
OPN3_4A .C..ACTCTTC...ATCA.CAC..CCA...TCTG..AC..G..C...........C....
OPN3_4B .C..ACTCTTC...ATCA.CAC..CCA...TCTG..AC..G..C...........C....
OPN3_4C .C..ACTCTTC...ATCA.CAC..CCA...TCTG..AC..G..C...........C....

OPN3_1  CTGTGGTCTCTCTACTAATAGCCTATGGGTATGGTCACCTTATAACTCCAACAGTAGCCA
```

```
OPN3_3  GGA.T..GG.G..GA.CGC.A.A.T....A.ACCCGGAT.A..T.....CT.T.CCAGT.
OPN3_4A GC..TA.AG.A..T..TGC.A.A.T...TCG.CC.GG.T.GG.GT....TGT..CCAGTG
OPN3_4B GC..TA.AG.A..T..TGC.A.A.T...TCG.CC.GG.T.GG.GT....TGT..CCAGTG
OPN3_4C GC..TA.AG.A..T..TGC.A.A.T...TCG.CC.GG.T.GG.GT....TGT..CCAGTG

OPN3_1  TTATCCCATCCTTCTTTGCCAAATCAAGCACTGCCTACAATCCAGTTATCTACATTTTCA
OPN3_3  .C..T..T...G.TC....A..GAGC.....C.T......C.....C..T.....C..TT
OPN3_4A .C..A..T..TA....G..A..GAGC..T....TA.T......CA.C.........C.A.
OPN3_4B .C..A..T..TA....G..A..GAGC..T....TA.T......CA.C.........C.A.
OPN3_4C .C..A..T..TA....G..A..GAGC..T....TA.T......CA.C.........C.A.

OPN3_1  TGAGCAGAAAGTTTCGCCGATGCCTTGTGCAGCTTTTTTGTGTGCAATTCCTGAGATTCA
OPN3_3  ...A..A.C.....TA.A.G...T..-..TGCTC..C.AAAA..TGG---.AAGA.A..C
OPN3_4A ...A..AGC....CTATAA....T..-CT..TGC..C.GCAC..-------.CAGCC...T
OPN3_4B ...A..AGC....CTATAA....T..-CT..TGC..C.GCAC..-------.CAGCC...T
OPN3_4C ...A..AGC....CTATAA....T..-CT..TGC..C.GCAC..-------.CAGCC...T

OPN3_1  AGAGGACCTTGAAAGAGCAACCGGCCATCGAGAGCAAC--AAACCTATCAGGCCTATTGT
OPN3_3  .TC.---...CC.GC.A...AT.TT.T..A..GT.G..TCG.GTG.G..G---.AGCATC
OPN3_4A TC..---T.GCTG.TG.AG.AACCAT..GCCAGT...----.GGTC..GG---.C..CCA
OPN3_4B TC..---T.GCTG.TG.AG.AACCAT..GCCAGT...----.GGTC..GG---.C..CCA
OPN3_4C TC..---T.GCTG.TG.AG.AACCAT..GCCAGT...----.GGTC..GG---.C..CCA

OPN3_1  CATGTCCCAGAAGGTAGGGGACAGGCCAAAGAAGAAAGTGACTTTTAGCTCTTCCTCAAT
OPN3_3  .GGAAA.AG..CAACTTCAC.TTT.TTGCT.CTTC.GCA.GACC--.C.--......GGA
OPN3_4A .CAAAA........C.CAA.GT--.GAGTTAT.TT..AAAG.CA--..TAG....A...A
OPN3_4B .CAAAA........C.CAA.GT--.GAGTTAT.TT..AAAG.CA--..TAG....A...A
OPN3_4C .CAAAA........C.CAA.GT--.GAGTTAT.TT..AAAG.CA--..TAG....A...A

OPN3_1  CATTTTCATAATCACCAGCGATGACACAGAGCAAATAGATGTCAGTACCAAATGTTCTGA
OPN3_3  -GCACCAGG.TG..ATTCTT.GC.TC....ATCC.G..CCCC.CACGGAC..CAGC.CC.
OPN3_4A TGGA.GAGA..G..ATCT---GTCTT.T...TCC.G..TCAAGCC.GGATCCG...T.AG
OPN3_4B TGGA.GAGA..G..ATCT---GTCTT.T...TCC.G..TCAAGCC.GGATCCG...T.AG
OPN3_4C TGGA.GAGA..G..ATCT---GTCTT.T...TCC.G..TCAAGCC.GGATCCG...T.AG
```

```
OPN3_1  CACAAAAATAAATGTGATCCAGGTGAAGC-CACTATAG-------
OPN3_3  AG-GC...AC..A.A.T.TTGCTC.T...T..T..C..TGTTTGA
OPN3_4A AGAGC.C.CCCCAACTG....A.GA..ATT..T.CCT.TGA----
OPN3_4B AGAGC.C.CCCCAACTG....A.GA..ATT..T.CCT.TGA----
OPN3_4C AGAGC.C.CCCCAACTG....A.GA..ATT..T.CCT.TGA----
;
end;
```

**Supplementary Table 2.**

Alignment of *Anolis carolinensis* coding sequences in the Opsin 4 gene family, including two paralogs (OPN4-1 and OPN4-2), and their predicted isoforms (2 isoforms of OPN4-1, 3 of OPN 4-2).

```
#NEXUS
[ Title NEXUS file]
begin data;
   dimensions ntax=5 nchar=1699;
   format    missing=?    gap=-    matchchar=.    datatype=nucleotide
interleave=yes;
   matrix

[!Domain=Data;]
OPN4_1A ------------------ATGGCCAACGCCTCGGAGGGGCCCCAGCAGATTCAGAGTATG
OPN4_1B ------------------------------------------------------------
OPN4_2A ATGTCTACAAGGCTGGTGGACTTGGCTC.A.T..TCCCAA.AG...G..CAAT.TTC.AC
OPN4_2B ATGTCTACAAGGCTGGTGGACTTGGCTC.A.T..TCCCAA.AG...G..CAAT.TTC.AC
OPN4_2C ATGTCTACAAGGCTGGTGGACTTGGCTC.A.T..TCCCAA.AG...G..CAAT.TTC.AC

OPN4_1A GGACCTCATCACAGAACAAAAGTGGATGTTCCTGACCATGTTCTTTACACCGTAGGTTCT
OPN4_1B ------------------------------------------------------------
OPN4_2A CTC..CG...CTTTTC...CG........A..AAC.....C..AC..T...A.T..GG..
OPN4_2B CTC..CG...CTTTTC...CG........A..AAC.....C..AC..T...A.T..GG..
OPN4_2C CTC..CG...CTTTTC...CG........A..AAC.....C..AC..T...A.T..GG..
```

```
OPN4_1A TGTGTTCTGGTAATCGGCTGTATTGGAATTACAGGAAACCTTCTCGTCCTTTATGCATTT
OPN4_1B ------------------------------------------------------------
OPN4_2A GTCA....T...G.G..AATC.CA....C.CTT..C........A...A.C....TT...
OPN4_2B GTCA....T...G.G..AATC.CA....C.CTT..C........A...A.C....TT...
OPN4_2C GTCA....T...G.G..AATC.CA....C.CTT..C........A...A.C....TT...

OPN4_1A TACAGCAACAAGAGGTTGAGGACTCCACCAAACTATTTCATAATGAATTTAGCAGTAAGC
OPN4_1B ------------------------------------------------------------
OPN4_2A .T...G.TTCG.G.CC.TC.......TG.C...ATG...G.T..C...C.G..T..C..T
OPN4_2B .T...G.TTCG.G.CC.TC.......TG.C...ATG...G.T..C...C.G..T..C..T
OPN4_2C .T...G.TTCG.G.CC.TC.......TG.C...ATG...G.T..C...C.G..T..C..T

OPN4_1A GATTTCCTGATGTCTGCAACTCAGGCTCCAATCTGTTTTCTCAACAGTATGCACAAGGAA
OPN4_1B ------------------------------------------------------------
OPN4_2A .....T..A...G.CATC......TG....G.T.TC..CAC......CC.CA....ACGG
OPN4_2B .....T..A...G.CATC......TG....G.T.TC..CAC......CC.CA....ACGG
OPN4_2C .....T..A...G.CATC......TG....G.T.TC..CAC......CC.CA....ACGG

OPN4_1A TGGGTACTTGGAGATATAGGTTGTAACTTGTATGCTTTTTGTGGGGCGCTCTTCGGAATA
OPN4_1B -AT.C.T..-----..C...........................................
OPN4_2A ...A..T....T.CA.A...C...G.GC....C..A..C.....A..T.....T..C..T
OPN4_2B ...A..T....T.CA.A...C...G.GC....C..A..C.....A..T.....T..C..T
OPN4_2C ...A..T....T.CA.A...C...G.GC....C..A..C.....A..T.....T..C..T

OPN4_1A ACCTCAATGATTACGTTGTTGGCTATTTCGGTTGATCGTTACTGTGTGATTACGAAGCCT
OPN4_1B ............................................................
OPN4_2A G.T..C.....C..C...ACA.TG..AG.CC.G..CA.G..T.T..........CGA..C
OPN4_2B G.T..C.....C..C...ACA.TG..AG.CC.G..CA.G..T.T..........CGA..C
OPN4_2C G.T..C.....C..C...ACA.TG..AG.CC.G..CA.G..T.T..........CGA..C

OPN4_1A CTCCAATCCATAAAAAGGACATCAAAAAAGCGCACCTGCATCATCATTGTCTTTGTCTGG
OPN4_1B ............................................................
OPN4_2A ..TGCC.....CGG.GCA.TG..T.CG...AAGG.T.TGC.A...C.ATCAGGA......
OPN4_2B ..TGCC.....CGG.GCA.TG..T.CG...AAGG.T.TGC.A...C.ATCAGGA......
```

```
OPN4_2C ..TGCC.....CGG.GCA.TG..T.CG...AAGG.T.TGC.A...C.ATCAGGA......

OPN4_1A CTCTACTCACTTGGATGGAGTGTATGTCCACTCTTTGGGTGGAGTTCCTATATACCAGAG
OPN4_1B ............................................................
OPN4_2A ..G..T..CT.A.CT......C.CCCG..TT.......A......G.T...G.T......
OPN4_2B ..G..T..CT.A.CT......C.CCCG..TT.......A......G.T...G.T......
OPN4_2C ..G..T..CT.A.CT......C.CCCG..TT.......A......G.T...G.T......

OPN4_1A GGTTTGATGATATCCTGTACATGGGACTATGTGACCTATTCTCCAGCAAATCGAAGTTAT
OPN4_1B ............................................................
OPN4_2A ..CC.CT.A.CC......T.C.....T...A.C..T.T.A.C...T..GTC..TGCC..C
OPN4_2B ..CC.CT.A.CC......T.C.....T...A.C..T.T.A.C...T..GTC..TGCC..C
OPN4_2C ..CC.CT.A.CC......T.C.....T...A.C..T.T.A.C...T..GTC..TGCC..C

OPN4_1A ACTATGATGTTGTGTTGCTGCGTGTTCTTTATCCCCCTCGTCATCATCTTCCACTGCTAT
OPN4_1B ............................................................
OPN4_2A ..G...C.CC.T.TC....TT..A........T.....AA.TGC....A.AT..A.T...
OPN4_2B ..G...C.CC.T.TC....TT..A........T.....AA.TGC....A.AT..A.T...
OPN4_2C ..G...C.CC.T.TC....TT..A........T.....AA.TGC....A.AT..A.T...

OPN4_1A ATATTCATGTTTCTGGCCATAAGAAGTACAGGCAGGAATGTTCAAAAACTGGGTTCAACC
OPN4_1B ............................................................
OPN4_2A G.G.....T..CA.A.....C.AG.A.T..AA....GC........GGACCAA....GA.
OPN4_2B G.G.....T..CA.A.....C.AG.A.T..AA....GC........GGACCAA....GA.
OPN4_2C G.G.....T..CA.A.....C.AG.A.T..AA....GC........GGACCAA....GA.

OPN4_1A TATAACCGGAAATCCTCTATTTCACACTCTA---TAAAGAGTGAATGGAAGCTTGCAAAA
OPN4_1B ...............................---..........................
OPN4_2A A.C.GTAAAG..GGTCAA.AACTCT..CAG.AGC.G....A...G.....AA.G..C...
OPN4_2B A.C.GTAAAG..GGTCAA.AACTCT..CAG.AGC.G....A...G.....AA.G..C...
OPN4_2C A.C.GTAAAG..GGTCAA.AACTCT..CAG.AGC.G....A...G.....AA.G..C...

OPN4_1A ATTGCCTTTGTGGCCATTGTGGTGTTCGTCTTGTCCTGGTCTCCATATGCTTGTGTTACT
OPN4_1B ............................................................
```

```
OPN4_2A G.A.....GA.T.T...CT..C.C.AT...A.C...............T..GTG..AG..
OPN4_2B G.A.....GA.T.T...CT..C.C.AT...A.C...............T..GTG..AG..
OPN4_2C G.A.....GA.T.T...CT..C.C.AT...A.C...............T..GTG..AG..

OPN4_1A CTAATTTCATGGGCAGGTTACGCCAGGACCTTAACTCCGTATTCTAAATCTGTGCCTGCT
OPN4_1B ............................................................
OPN4_2A T.GG.GG.T.TT.....A...T.TCATCT.C.T..A..A..CATG..C..A..C.....A
OPN4_2B T.GG.GG.T.TT.....A...T.TCATCT.C.T..A..A..CATG..C..A..C.....A
OPN4_2C T.GG.GG.T.TT.....A...T.TCATCT.C.T..A..A..CATG..C..A..C.....A

OPN4_1A GTTATTGCCAAAGCTTCTGCAATCTACAATCCCATAATCTATGCAATCATTCACCCAAGA
OPN4_1B ............................................................
OPN4_2A ..G........G.......TT...C....C..A..C..T.....C..TG....T..C.A.
OPN4_2B ..G........G.......TT...C....C..A..C..T.....C..TG....T..C.A.
OPN4_2C ..G........G.......TT...C....C..A..C..T.....C..TG....T..C.A.

OPN4_1A TACAGAAGAACCATTCGAAGTGCTGTTCCTTGCTTGAGATTTCTTATACCGATATCTAAA
OPN4_1B ............................................................
OPN4_2A .......TGG.G...GC..AGTT.C....A...C.TG...C...GC..AGAG.TC.CCGC
OPN4_2B .......TGG.G...GC..AGTT.C....A...C.TG...C...GC..AGAG.TC.CCGC
OPN4_2C .......TGG.G...GC..AGTT.C....A...C.TG...C...GC..AGAG.TC.CCGC

OPN4_1A AGTGATCTTTCAACAAGTTCTATGAGTGAGTCATCGTTCCGGGCTTCTGTATCTAGCCGT
OPN4_1B ............................................................
OPN4_2A .AG..C--.C..G.T.CCC..CCACCA..-----...C..A-----....CA.C....AG
OPN4_2B .AG..C--.C..G.T.CCC..CCACCA..-----...C..A-----....CA.C....AG
OPN4_2C .AG..C--.C..G.T.CCC..CCACCA..-----...C..A-----....CA.C....AG

OPN4_1A CACTCT--TTCAGCTACAGGAACAAGAGCACTTACA----TTTCCTCCATTTCTGCTAAA
OPN4_1B ......--............................----....................
OPN4_2A TC....GA.AT.AA.GG..TGC......G..ACCG.AGGT.A.....TG.C..A.AC.GT
OPN4_2B TC....GA.AT.AA.GG..TGC......G..ACCG.AGGT.A.....TG.C..A.AC.GT
OPN4_2C TC....GA.AT.AA.GG..TGC......G..ACCG.AGGT.A.....TG.C..A.AC.GT
```

```
OPN4_1A GAAACAACTTGGTGTGATGTAGAGCTTGACCCTGTTGAGTCAGGCCACAAGAAGTTGCAA
OPN4_1B ............................................................
OPN4_2A ...T.TGA....AC...CAC...AGC...TATCAG.-..C...AATT.C...-..C..C.
OPN4_2B ...T.TGA....AC...CAC...AGC...TATCAG.-..C...AATT.C...-..C..C.
OPN4_2C ...T.TGA....AC...CAC...AGC...TATCAG.-..C...AATT.C...-..C..C.

OPN4_1A GCCTACCGAAGTAATTCTTTTTCAGCCAAAGGAGTAGCAGAAGAAGAGAGTGGGTTGCTT
OPN4_1B ............................................................
OPN4_2A .TGGGT.A.T----...C.ACAG.ATAT.T.A..ACA..ACT...ACC.TCAA.G.CAAA
OPN4_2B .TGGGT.A.T----...C.ACAG.ATAT.T.A..ACA..ACT...ACC.TCAA.G.CAAA
OPN4_2C .TGGGT.A.T----...C.ACAG.ATAT.T.A..ACA..ACT...ACC.TCAA.G.CAAA

OPN4_1A CTCAGAACAAATAACTGCAATGTGCCAGCAAGAAAAAAGGTTGCTCTGTCATCCATCAGT
OPN4_1B ............................................................
OPN4_2A TCT.A..TG.GA.G.CATG..TCAGGGATTTTTG...G.---------------------
OPN4_2B TCT.A..TG.GA.G.CATG..TCAGGGATTTTTG...G.AC.T.CG..GATG..GATGAC
OPN4_2C TCT.A..TG.GA.G.CATG..TCAGGGATTTTTG...G.---------------------

OPN4_1A CTAGAAGAATCACATGCACAATCCTTTGGAGGGAACAGCTCAGAACTGTTGCTCCTAAGC
OPN4_1B ............................................................
OPN4_2A ------------------------------------------------------AC.G.T
OPN4_2B A.TTCTATGGTGG.G.ACTT.C.........TCT.T.A.CTT...GATAGTT.GAC.G.T
OPN4_2C ---------------.ACTT.C.........TCT.T.A.CTT...GATAGTT.GAC.G.T

OPN4_1A AGTTCCTTGAAAACATCTTCTCTTCCATTTGGCTTGAATGGTAGCAGCACAGGAGAAAGT
OPN4_1B ............................................................
OPN4_2A GAAGATC....TG.C.-----------------...G....-..A..GGAG.AGTCTTA.
OPN4_2B GAAGATC....TG.C.-----------------...G....-..A..GGAG.AGTCTTA.
OPN4_2C GAAGATC....TG.C.-----------------...G....-..A..GGAG.AGTCTTA.

OPN4_1A ACTGATACTTCTCAATTGGAAGGCCAAGAAAACCAAATAAATGGAAGCCTGGACTCCTTC
OPN4_1B ............................................................
OPN4_2A T.C.GGC.A...ACT.C---.A.....ATCCC.AGC..C.T..TG.C.T.CAG.AATG..
OPN4_2B T.C.GGC.A...ACT.C---.A.....ATCCC.AGC..C.T..TG.C.T.CAG.AATG..
```

```
OPN4_2C T.C.GGC.A...ACT.C---.A.....ATCCC.AGC..C.T..TG.C.T.CAG.AATG..

OPN4_1A ACAAGCCCTATTCTTCCACAAATTATTATAATTCCTACCTCAGAGACCAAT-CTCTCTGA
OPN4_1B ...................................................-........
OPN4_2A CAG..AA.AGAC..G..--------..TGG.G...AG.TCTG.T.C.TTG.G...CAG..
OPN4_2B CAG..AA.AGAC..G..--------..TGG.G...AG.TCTG.T.C.TTG.G...CAG..
OPN4_2C CAG..AA.AGAC..G..--------..TGG.G...AG.TCTG.T.C.TTG.G...CAG..

OPN4_1A GGGGCAAGCAGAACCAGAAAATGCACAAGATGAAAATATAGATTTATTTTTTTCTCAAGG
OPN4_1B ............................................................
OPN4_2A ACA.T.GCT.T.G.TG.G...A.G.TT---CC..C.GCCGTC..G.A.GG..ATG.C.T.
OPN4_2B ACA.T.GCT.T.G.TG.G...A.G.TT---CC..C.GCCGTC..G.A.GG..ATG.C.T.
OPN4_2C ACA.T.GCT.T.G.TG.G...A.G.TT---CC..C.GCCGTC..G.A.GG..ATG.C.T.

OPN4_1A GAAGAACCACCTCCTGGACATAGGAAGGCTTAGTTCCTCTACTGAACTCCTTGAAGCAAT
OPN4_1B ............................................................
OPN4_2A CG..TG..TTAG.TCCT..C.CT...A.AAATA.GTG..AG..C.G.G--.A....ATC.
OPN4_2B CG..TG..TTAG.TCCT..C.CT...A.AAATA.GTG..AG..C.G.G--.A....ATC.
OPN4_2C CG..TG..TTAG.TCCT..C.CT...A.AAATA.GTG..AG..C.G.G--.A....ATC.

OPN4_1A TGAGAAGCTTCTGTCATAA
OPN4_1B ...................
OPN4_2A ..CA.GA.AGG.T.A.---
OPN4_2B ..CA.GA.AGG.T.A.---
OPN4_2C ..CA.GA.AGG.T.A.---
;
end;
```

**Supplementary Table 3.** Alignment of *Anolis carolinensis* coding sequences in the Opsin 5 gene family, including four paralogs (OPN5-1, OPN5-2, OPN5-4, and OPN5-6), and three predicted isoforms of OPN5-4.

```
#NEXUS
[ Title Opsin 5]
```

```
begin data;
   dimensions ntax=6 nchar=1390;
   format    missing=?    gap=-    matchchar=.    datatype=nucleotide
interleave=yes;
   matrix

[!Domain=Data property=Coding CodonStart=1;]
OPN5_1  ATGGAACAAGGCCAAAATATCTCATCTCAAGATGATAACCAGCAAGAGGAAGACCCCTTT
OPN5_2  -------------.TGTCTCTC..GG.TTCTCCCCAGG.ACCATG...C..TGT.A....
OPN5_4A -------------------------------...GAGT.TTA.TTTGCA..T..GA.A..
OPN5_4B -------------------------------...GAGT.TTA.TTTGCA..T..GA.A..
OPN5_4C -------------------------------...GAGT.TTA.TTTGCA..T..GA.A..
OPN5_6  ------------------------------------------ATG.....GC..TA.A.C

OPN5_1  GCGT-CAAAAC---TATCAGTTGAAGCAGACATAGTAGCTGGAGTTTATTTATTAGTAAT
OPN5_2  TTCAAAC...GAGG.CC.T..CTCG.G...G.CTA.TAT...CT.C...C.TC.G.CTC.
OPN5_4A T.A.T.C...A---..A...AA.C...G..TG.TA.T.T...C..C.T..ATA.T...T.
OPN5_4B T.A.T.C...A---..A...AA.C...G..TG.TA.T.T...C..C.T..ATA.T...T.
OPN5_4C T.A.T.C...A---..A...AA.C...G..TG.TA.T.T...C..C.T..ATA.T...T.
OPN5_6  T.----C...G---.TCACCCA.T.TGG..TTAT.G.ATG..T..G.T.C.TC.GA.T..

OPN5_1  TGGGATTTTGTCAACTTTAGGAAATGGATACGTTATTTACATGTCCACCCAACGGAAAAA
OPN5_2  A..TTGGA....TTGG..T..C..CA.CATT..A..C.TTG.ACTATATAG..A.CG.GC
OPN5_4A ...C..A.GC..TTT..GT..G..CA.TAT.C.CC.C..TG.T...TA.A.GAAA..G..
OPN5_4B ...C..A.GC..TTT..GT..G..CA.TAT.C.CC.C..TG.T...TA.A.GAAA..G..
OPN5_4C ...C..A.GC..TTT..GT..G..CA.TAT.C.CC.C..TG.T...TA.A.GAAA..G..
OPN5_6  ..CA..C...A...T.C.G..G...TCGATG...C.AGCAG.AG.TGTGA.G...TCCTC

OPN5_1  GAAGCTGAAGCCTGCTGAAATAATGACTGTCAATTTAGCGGTGTGTGACTTGGGAATTTC
OPN5_2  TGGA..TC....GA.A..TTACC....AT.T..CC.G..T..A.C...TGCCA.TG.C..
OPN5_4A CCTAT....A..A..A...TACT.C.TGA....C..G..CA.CA....TC.T..T..GA.
OPN5_4B CCTAT....A..A..A...TACT.C.TGA....C..G..CA.CA....TC.T..T..GA.
OPN5_4C CCTAT....A..A..A...TACT.C.TGA....C..G..CA.CA....TC.T..T..GA.
OPN5_6  CTGC..C.G.T.AC....GC.CC.C..A..........A.CCAC...TC.T.....GGG
```

```
OPN5_1  AGTGGTGGGAAAGCCTTTTAGCATCAT-----TGCCTTCTTCTCCCACCGCTG---GATT
OPN5_2  T..TT.T..TT.TT.CCGGG.G..A..AGAAA.TTTCAA.GTATT..GA.A..ATG....
OPN5_4A TC..ACCTTGT.C...C.AGCTG.T.C-----AT..AG.C.TG.A...A.G..---.C.G
OPN5_4B TC..ACCTTGT.C...C.AGCTG.T.C-----AT..AG.C.TG.A...A.G..---.C.G
OPN5_4C TC..ACCTTGT.C...C.AGCTG.T.C-----AT..AG.C.TG.A...A.G..---.C.G
OPN5_6  TC.CAGCATGT.T..A..GGC...TGC-----.T.TGC..GGAA....GCT..---.C.A

OPN5_1  TTTGGTTG-------GAGTGGTTGCCGTTGGTATGGATGGGCTGGGTTCTTCTTTGGCAT
OPN5_2  CC.TA.CACATCGATAT.GAC.....AGGT.G......TTCTGAC.C.AC.G.....TC.
OPN5_4A .....GCA-------ACAA.T...TTTA.TC....C..TTTG...AG.G..T.....AG.
OPN5_4B .....GCA-------ACAA.T...TTTA.TC....C..TTTG...AG.G..T.....AG.
OPN5_4C .....GCA-------ACAA.T...TTTA.TC....C..TTTG...AG.G..T.....AG.
OPN5_6  GG...GGA-------.GCAACC...AT..AC....CTCT.ATG..C..TC.T..C..TG.

OPN5_1  TGGGAGTCTTATTACCATGACAGCAGTAAGCTTGGATCGATATTTCAAAATCTGCCATTT
OPN5_2  ..CA..CA...A...AC......TGA.T...G..ACC......A.A...GGA......CC
OPN5_4A CT.C.....A.C...GC.....CT.C.G..TA.A.TGT.T.GCC.A.....T...TT.CC
OPN5_4B CT.C.....A.C...GC.....CT.C.G..TA.A.TGT.T.GCC.A.....T...TT.CC
OPN5_4C CT.C.....A.C...GC.....CT.C.G..TA.A.TGT.T.GCC.A.....T...TT.CC
OPN5_6  CTCC..CA.A..G..TT..T.T...A.GGCTG.AATC..T.TCC.TGT..CT.TTTCA.C

OPN5_1  AT--CCTATGGTACCTGGCTGAAGAGACACCATGTT-TTTATCTGCTTAGGAATCATCTG
OPN5_2  TG---ACCGA.G..AGT.TATC..CA.T.G..GCA.C.C.G.GGCTC.TTTTC.G..A..
OPN5_4A .G--TT.....G.A.A.AT.T.GACC.GGA....G.-.GG..A.TGA...CCTGTGCT..
OPN5_4B .G--TT.....G.A.A.AT.T.GACC.GGA....G.-.GG..A.TGA...CCTGTGCT..
OPN5_4C .G--TT.....G.A.A.AT.T.GACC.GGA....G.-.GG..A.TGA...CCTGTGCT..
OPN5_6  .AAA...GC.AGG.A.AAAATC.ACAGG.AAG..A.GCA...T..TA..ATGC.G..T..

OPN5_1  GTCCTATGCTGCCTTCTGGGCAACCATCCCTTTTGCAGGCTTTGGCAATTATGCTCCTGA
OPN5_2  .ATTGCA...TT.......T..GTAGCT..AG.CCTG..A.GG....G....A.AGA.CG
OPN5_4A .GT...C..A...A.T.TT..CTTTTC...AC.A..TCA..GG..AG.G....GAG.A..
OPN5_4B .GT...C..A...A.T.TT..CTTTTC...AC.A..TCA..GG..AG.G....GAG.A..
OPN5_4C .GT...C..A...A.T.TT..CTTTTC...AC.A..TCA..GG..AG.G....GAG.A..
```

```
OPN5_6  .G.A.....A.T.C.T.....C.TTC.G.....GCTG....GG..AC......GC..A..

OPN5_1  GCCATTTGGGACCTCGTGCACTCTGGACTGGTGGCTGGCTCAAGGTTCAGTGGCTGGGCA
OPN5_2  CATG.A...A..T---..TGAAA.C......GCAAAA..C-..CT.CTCAACAA.CTA..
OPN5_4A ...T.A...T..AG.A..TTGCA.T..T...C.TA.AT.A-..CA.GA..AA.ACA.C..
OPN5_4B ...T.A...T..AG.A..TTGCA.T..T...C.TA.AT.A-..CA.GA..AA.ACA.C..
OPN5_4C ...T.A...T..AG.A..TTGCA.T..T...C.TA.AT.A-..CA.GA..AA.ACA.C..
OPN5_6  ......C..C..T..T......A.C.CA...G.C.A.TTC..CAA...GCAAAAG..ATT

OPN5_1  GGCCTT-TATTCTGAATATCCTTTTCTTCTGTCTTGTGCTTCCAACTGCTGTGATTATGT
OPN5_2  A.T...A....G..TCC..AT.CA...GT...T..T.C..A..TGTGT....C..GG.T.
OPN5_4A T.T...AC.C.ACTGCCC..T..G.......CTACA.CA.C..CTG..GAA.A.....TA
OPN5_4B T.T...AC.C.ACTGCCC..T..G.......CTACA.CA.C..CTG..GAA.A.....TA
OPN5_4C T.T...AC.C.ACTGCCC..T..G.......CTACA.CA.C..CTG..GAA.A.....TA
OPN5_6  T.....-C.......GC..GT.CA.TC....CACAT.T..G...G.AATAACC..C..CA

OPN5_1  TTTGCTATGTTAAAATTATTGCAAAAGTACAGT-CATCCACCAAAGAAGTAGCCCACTAT
OPN5_2  ...C......GTCC..C...AAT.C...CA...CA.G....GCTCT...CG.AGTGGGTG
OPN5_4A C..CT..CAC.CTT..CC.GATT.CT..GA..G-AT....GA....C...G.AA..AC..
OPN5_4B C..CT..CAC.CTT..CC.GATT.CT..GA..G-AT....GA....C...G.AA..AC..
OPN5_4C C..CT..CAC.CTT..CC.GATT.CT..GA..G-AT....GA....C...G.AA..AC..
OPN5_6  .G..T..CT.GGG...CGCCTGG...T.T..TA-A.A.TCA.C.....A.GCAAA...TA

OPN5_1  GACACCAGGATTCAAAACCAACATGTACTGGAAATGAAGTTA--ACTAAGGTGGCAATGT
OPN5_2  AC.----CA.CGG.G.GA-C.G.G.CGTAT.G..C.....-GTC..A.G...TT.TG.TG
OPN5_4A .C.-TTG...CC..CC.G-G.TGA.C.G..TGC.CACCA...CAG.C...C..AGC--A.
OPN5_4B .C.-TTG...CC..CC.G-G.TGA.C.G..TGC.CACCA...CAG.C...A..CTCT.A.
OPN5_4C .C.-TTG...CC..CC.G-G.TGA.C.G..TGC.CACCA...CAG.C.-----AGC--A.
OPN5_6  A..---..A..ATC..GTGC.GCAAAG.....G.A......GAT...G--.....TG.A.

OPN5_1  TGATCTGTGCA--GGATTTATGTTTGCTTGGATCCCATATGCCGTGGTGTCTGTGTGGTC
OPN5_2  .......CA.T--.CC.....CACG..C...TCA..G.....T..CA.C..CA.......
OPN5_4A C.C.G.A....TT......T.TG.G..C....GT........TA.CA.TG.AA.....G.
OPN5_4B ....ACT.A..TCA.CA..TCAGCAAAAATA.----------------------------
```

```
OPN5_4C .TCAGCAAAA.TAAA.GC.GA---------------------------------------
OPN5_6  .....A...T.--..T..CC.TGGC..A....C.........TA.A..CAGCT.C.....

OPN5_1  AGCTTTCGGAAGACCAGATTCCGTTCCCATCAAAGTTTCAGTGATACCAACTTTGCTTGC
OPN5_2  T..C.AT..TT------.CA.T..G....ATTT.AC.AGCA.CC.TG.C.GCC.TT....
OPN5_4A C..C..T...TC.ATT..CATGA....TCCACT..C..TT.CAG.T...G..G.AT....
OPN5_4B ------------------------------------------------------------
OPN5_4C ------------------------------------------------------------
OPN5_6  ..TA..TCACTCCAGT..A..AA.A..TTA..TT...A.GT.AT....TTG.C..T....

OPN5_1  AAAATCAGCAGCCATGTACAACCCAGTTATTTACCAGGTCATTGACTGCAAATCTGCCTG
OPN5_2  G..G..T..CAG.T.T.....T..CA.C......TTT.GA..GAG..C.....TCCGAAA
OPN5_4A ...G..TT..A.AC.T........G.C...G..T.TTT.TC.AA.GCC...T.TCCGAA.
OPN5_4B ------------------------------------------------------------
OPN5_4C ------------------------------------------------------------
OPN5_6  T..G...T..A.AGCC.....T..TT.C..A...T.CACTT.CAG.AAA.C..TCCGGCA

OPN5_1  TTGCCGGCCAGGTAACCTCCAGCCACTGCAGAAGAAGAACTCCAGGCTGTACATAATACC
OPN5_2  G----.A.ATCT.CGT..TGCT......TGCC.A.GAG.TCAAG.ATCC.GTGA.GCT.A
OPN5_4A CAC.ATTG.CAAAG.T..TACAGT...C..CCGATTATGTCT..AAAGC.G.T.TTGT..
OPN5_4B ------------------------------------------------------------
OPN5_4C ------------------------------------------------------------
OPN5_6  .-.AA.TTA..CATCT.AGAT.TT.T.CTG.CCA....G...A..AAAAC.TGA..A..T

OPN5_1  TACAGGAAAAAAATCTGAGGTAGTTCAGGAAACACAACTGGACAGTGTATGA--------
OPN5_2  A..G.TTC..G..C..C.A.C..AAACA.G..GTTTCACCTT.TCAAAGA..AGAAAAAT
OPN5_4A C.G...C.TGC.GAACTGCTCTTACAGATC.G.T.TTGA..CTCCAC.GAAGTCATTTAA
OPN5_4B ------------------------------------------------------------
OPN5_4C ------------------------------------------------------------
OPN5_6  CCATCA.T.GC...G.ATCA.TCA.GT..C.CGGAGGAG.C.ATA.C.G.CTTTCAACCC

OPN5_1  ------------------------------------------------------------
OPN5_2  ACGCTGCAGATGTCCAGCCTGCTCTAAGCCCTGATTCTGGGGTAGGGAGTCGTTCAAACA
OPN5_4A AGGAAGAAATGAATCAAGCAGTAACTCTGTGCAAATCGTGGGAGGATGCTCATACTTTCC
```

```
OPN5_4B ------------------------------------------------------------
OPN5_4C ------------------------------------------------------------
OPN5_6  GACAGATAGAAATGAGAGAAATACCAAATCAATAG-------------------------

OPN5_1  ------------------------------------------------------------
OPN5_2  CTCCTCCCCCAGTAAACAGGGAAGTGTATTTCGGTGCTTTTGATACCTTTTCCAACAATC
OPN5_4A TTGTGAGAAGTGCCATGATCCCTTTGAATGCTTCAAGAACTATCCAAAATGCTGCCAAGG
OPN5_4B ------------------------------------------------------------
OPN5_4C ------------------------------------------------------------
OPN5_6  ------------------------------------------------------------

OPN5_1  ------------------------------------------------------------
OPN5_2  CTGATGTTGAATGTGACAGACTGTGA----------------------------------
OPN5_4A GAGGCTAAATGTTATGGATCATACTCCTCGAGAAAGTATATCAGTGGAAAATAACATGCA
OPN5_4B ------------------------------------------------------------
OPN5_4C ------------------------------------------------------------
OPN5_6  ------------------------------------------------------------

OPN5_1  ------------------------------------------------------------
OPN5_2  ------------------------------------------------------------
OPN5_4A ATCCAAAACAAAACATGCTTCTGAAAAATATATAAAGGTTGTGATCAGGGGAGAAAAAAA
OPN5_4B ------------------------------------------------------------
OPN5_4C ------------------------------------------------------------
OPN5_6  ------------------------------------------------------------

OPN5_1  ------------------------------------------------------------
OPN5_2  ------------------------------------------------------------
OPN5_4A CACTGATATTGATAATTTGGAAATAACATTAGAACACATACCTACAGATATTAAATTTGC
OPN5_4B ------------------------------------------------------------
OPN5_4C ------------------------------------------------------------
OPN5_6  ------------------------------------------------------------

OPN5_1  ----------
OPN5_2  ----------
```

```
OPN5_4A CAATCTTTAG
OPN5_4B ----------
OPN5_4C ----------
OPN5_6  ----------
;
end;
```

**Supplementary Table 4.** Primers for green anole (Anolis carolinensis) opsin transcripts

| Primer Name | Sequence |
|---|---|
| **Opn1VA_1F** | ATGGCTGGACTGAGGAGGG |
| **Opn1VA_1R** | TTTCGGAAATTGCCAGCTGAG |
| **Opsin31_1F** | ATGTTCTCCGCCAACG |
| **Opsin31_1R** | CTATAGTGGCTTCACCTGGATC |
| **Opsin33_F** | GTAGCTATGGACCTGAAGGAC |
| **Opsin33_R** | TCAAACACTGTAATGAGCTACGAG |
| **Opsin3-4a_1F** | ATGTCAGAACTTAGCTCCAATTTG |
| **Opsin3-4a_1R** | TCACAGGAATGAATTTTC |
| **Opsin4-1a_1F** | ATGGCCAACGCCTCGGAG |
| **Opsin4-1a_1R** | TTATGACAGAAGCTTCTCAATTG |
| **Opsin4-1a_2F** | GCCCCAGCAGATTCAGAGTAT |
| **Opsin4-1a_2R** | GGTTCTGCTTGCCCCTC |
| **Opsin42a_F** | ATGTCTACAAGGCTGGTGGA |
| **Opsin42a_R** | TTAAACCTGTCTTGCAAGATCTTCTACGC |
| **Opsin5-2_1F** | ATGTCTCTCCAGGTTTCTCCCC |
| **Opsin5-2_1R** | TCACAGTCTGTCACATTCAACATC |
| **Opsin5-6_1F** | ATGGAGGAGCACTACATCTC |
| **Opsin5-6_1R** | CTATTGATTTGGTATTTCTCTCATTTCTATCTG |

**Supplementary Figure 1**. Gene tree of *Anolis carolinensis* opsin coding sequences, constructed in MEGA-X using Maximum Likelihood and the Tamura-Nei model. Bootstrap values are indicated by each node.

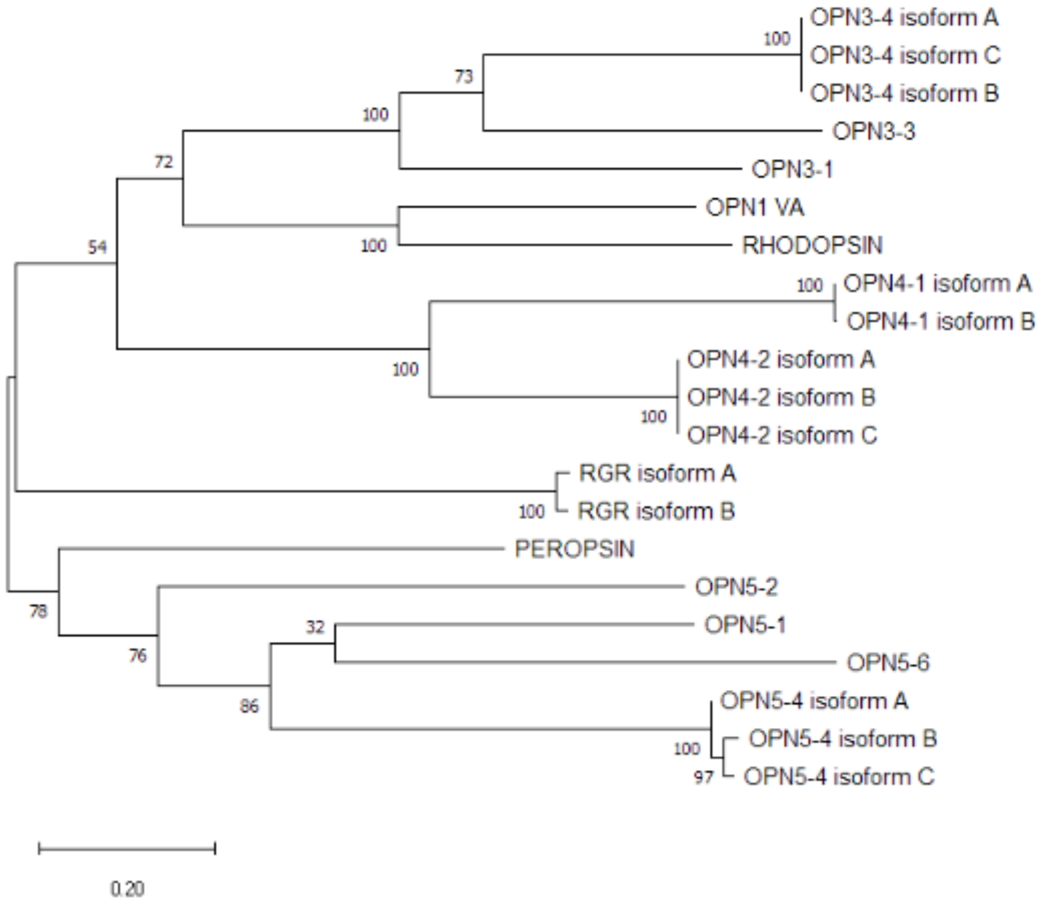
